# Exploring the Antidepressant Effects of Saffron Constituents: Targeting Dopamine and Serotonin Transport Proteins, and Monoamine Oxidase-B: An in Silico Evidence-Based Study

**DOI:** 10.64898/2026.03.16.712249

**Authors:** Brijendra Singh, Deepak Sharma, Vyas Madhavrao Shingatgeri, Vinay Lomash

## Abstract

Globally, about 264 million individuals across all age groups are impacted by depression, a prevalent central nervous system (CNS) condition. Chronic and enduring depression might result in significant health consequences. Numerous pharmaceutical antidepressants exist for the management of mild to severe depression, largely functioning by modifying neurotransmitter levels in the brain. Nevertheless, these drugs frequently induce a variety of side effects, such as insomnia, constipation, exhaustion, drowsiness, and anxiety. Saffron (*Crocus sativus L*.) is widely acknowledged as a natural antidepressant with little adverse effects. This study investigated the potential antidepressant mechanisms of saffron’s principal bioactive compounds safranal, crocin, and picrocrocin via molecular docking against critical target proteins associated with depression, namely the dopamine transporter (DAT), serotonin transporter (SERT), and monoamine oxidase B (MAO-B). Molecular docking was conducted with AutoDock 4.2 to assess the binding affinity and interaction energy of these drugs with the target proteins. Furthermore, Discovery Studio facilitated the viewing and study of both interacting and non-interacting residues at the docking sites, juxtaposing these interactions with those of established inhibitors in crystal structures. The permeability of the blood-brain barrier (BBB), pharmacokinetic characteristics, and toxicity profiles of saffron components were evaluated using SWISS ADME, DataWarrior, and Osiris Molecular Property Explorer. Among the evaluated elements, safranal had the greatest potential as a competitive inhibitor of the dopamine transporter, according to its notable blood-brain barrier permeability, robust binding affinity, and analogous interaction residues in comparison to nortriptyline, a recognized inhibitor. Our findings indicate that safranal may be a viable natural alternative to traditional antidepressants, with minimized adverse effects.

## 1. Introduction

Saffron is a widely used spice derived from the blooms of *Crocus sativus L*. (*C. sativus*), which belongs to the Iridaceae family. *Crocus sativus L*., used to produce saffron, was first planted in Greece. However, countries such as Iran, Turkey, India (Kashmir), Afghanistan, and Spain are currently regarded as major cultivators of *Crocus sativus L*. (Tammaro, 1990; Srivastava et al., 2010). Saffron is a threadlike crimson stigma from the plant *Crocus sativus L*. that is used as a spice to impart unusual aroma and color to food preparations and other items. Saffron is also recognized as a folk medicinal herb that can treat various illnesses (Hosseinzadeh, 2014; Razak et al., 2017). Saffron has been shown to have anticonvulsant properties (Menghani et al., 2021), regulate neurotransmitter levels like dopamine and glutamate (Ojetunde, 2024), improve learning and memory, have anti-Alzheimer’s and anti-Parkinson’s, be anxiolytic and hypnotic, reduce morphine dependency, and protect against brain ischemia (Khazdair et al., 2015; Abdian et al., 2024). Furthermore, it is used medicinally to treat genital and urinary system diseases such as erectile dysfunction and ejaculatory latency (Najafabadi et al., 2022). As a result, modern research is increasingly focused on analytical investigations for extracting and purifying saffron components for pharmacological and therapeutic applications. Chemical investigation of *C. sativus* revealed more than 150 chemicals in the saffron stigma (Winterhalter and Straubinger, 2000). Chemical ingredients are divided into volatile and non-volatile components. More than 34 volatile components are detected, including terpenes, terpene alcohols, terpene esters, and safranal (Liakopoulou-Kyriakides, 2002). The interaction of heat and light during the drying process produces safranal, the primary volatile component of picrocrocin (Avila-Sosa et al., 2022). Non-volatile components include crocin, -crocin, α crocetin, picrocrocin, and carotenoids (zeaxanthin, lycopene, and α -carotene). Crocetin and crocins are colored carotenoids with glycosidic derivatives (Wallis, 1958; Akowuah and Htar, 2014). According to Conrad and Yen (2007), non-glycosylated carotenoids include - β carotene, lycopene, and zea-xanthin (Conrad, 2007). Crocin (C44H64O24), picrocrocin (C16H26O7), and safranal (C10H14O) are the principal bioactive chemical components of saffron (Khorasany and Hosseinzadeh, 2016). These primary components play a crucial part in saffron’s therapeutic effects on various organs of the body. The blood-brain barrier (BBB) is a significant impediment to treating neurodegenerative illnesses. To combat neurodegenerative etiology, drugs should be delivered to the central nervous system (CNS) after crossing the blood-brain barrier and traversing to come together with its corresponding target in the brain adenexa to mimic the concert of chemical changes necessary for depicting its biological activity (Niazi, 2023). In this study, natural bioactive compounds (picrocrocin, safranal, and crocin) found in saffron were first tested for blood-brain barrier permeability, and then, additionally, these molecules were docked to their target protein. To establish the biological activity of every newly created chemical or its An analogy: The literature contains a vast number of animal models for diverse diseases. Similarly, numerous studies demonstrate the biological efficacy of drugs against neurodegenerative disorders such as Parkinsonism, Alzheimer’s, and multiple sclerosis. However, therapeutic molecule screening for each condition requires many animals and extensive chemical synthesis, which is neither cost-effective nor ethical. Recently, machine learning approaches in computer science have resulted in powerful software that can precisely predict the simulated chemical, physical, and biological characteristics of pharmaceutical compounds. Predicting the chemical, physical, and biological activity of a drug candidate using such software is known as "in-silico analysis," and it helps to reduce the usage of animals, as per the principle of 3Rs (Replacement, Reduction, and Refinement), and save time and resources. Additionally, we use molecular docking to screen the activity of pharmacological compounds against recognized receptors that regulate the synthesis and inhibition of specific neurotransmitters. This in-silico study estimates how well the potential saffron components work and their potential harm. The most promising component can then be tested and validated by in vivo research using a preferred animal model to confirm effectiveness and toxicity, which also helps to minimize the use of animals. The current study aimed to conduct an in-silico analysis to screen the natural bioactive elements of saffron for their ability to cross the blood-brain barrier (BBB) after being absorbed through the stomach and delivered to the central nervous system (CNS). Furthermore, molecular docking was performed using AutoDock software to determine the optimal binding posture and rank it based on binding affinity, as well as to forecast the stability of each individual pose with dopamine transporter, serotonin transporter, and monoamine oxidase B receptors.

## 2. Materials and methods

Biological activity of bioactive components of saffron (picrocrocin, safranal, and crocin) was evaluated through in silico analysis methods, utilizing computer-based techniques like molecular docking and toxicity prediction to effectively screen and assess potential therapeutic candidates.

### 2.1 In silico analysis methods

This study utilized the SwissADME web tool to perform *in silico* biological activity analysis of the synthesized molecules. SwissADME (http://www.swissadme.ch) is a freely accessible, user-friendly interface that can be employed by both specialists and non-experts in cheminformatics or computational chemistry to predict key pharmacokinetic parameters and support drug discovery efforts. In this study, we specifically assessed parameters related to gastrointestinal absorption and blood–brain barrier (BBB) permeability. SwissADME offers strong prediction tools for how drugs are absorbed in the body and how easily they can cross the blood-brain barrier, using its methods like the BOILED-Egg model, iLOGP, and the Bioavailability Radar. This tool yielded predictions for gastrointestinal absorbability and BBB permeability, as shown in Figures 1 and 2, respectively.

**Fig. 1.**
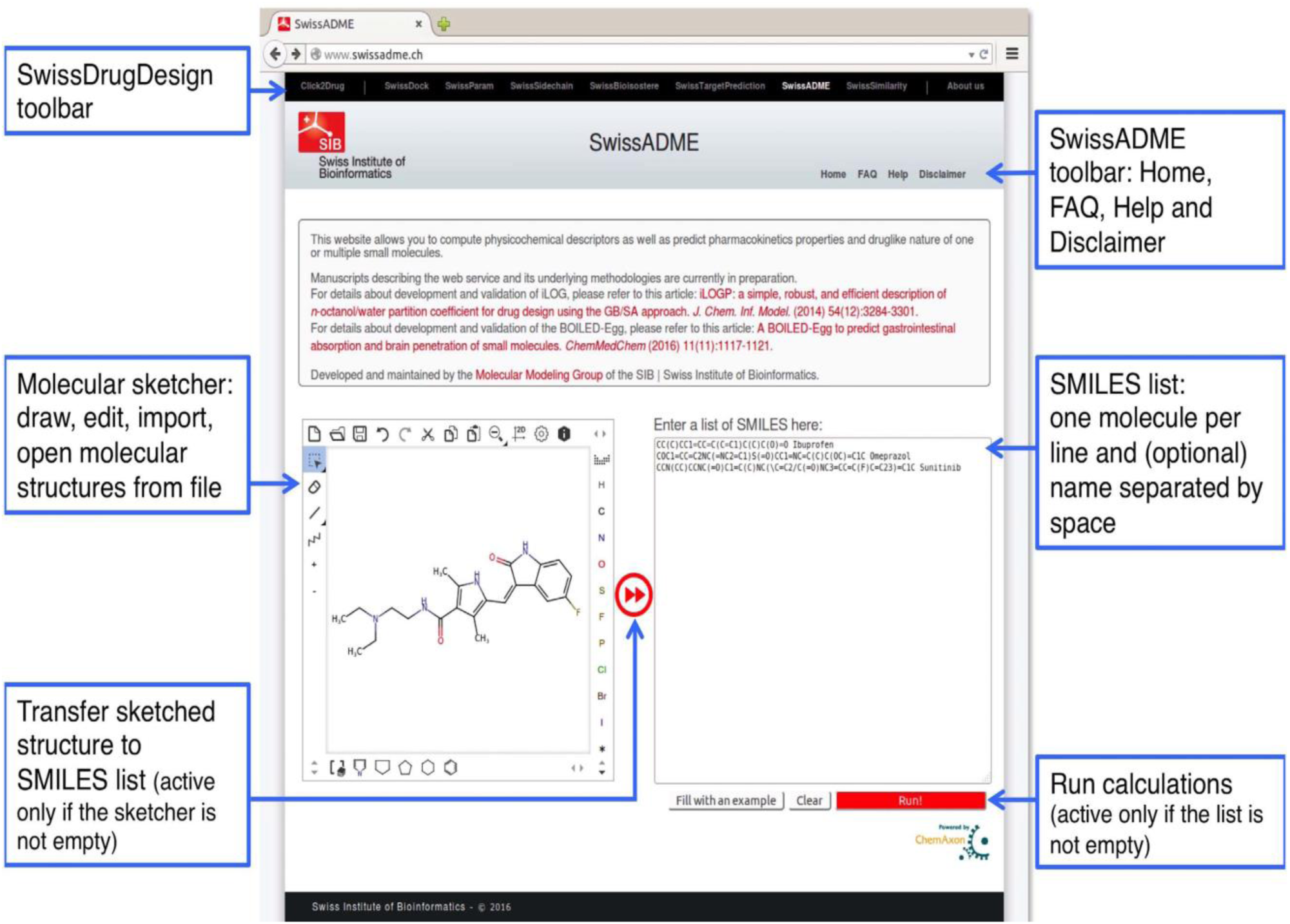
Swiss ADME submission page. **Structure of** Molecules can be pasted or drawn with the help of molecular sketcher. Further process as shown in the flow chart in the figure. Lastly user can start the calculations by clicking on the “Run” button.

**Fig. 2.**
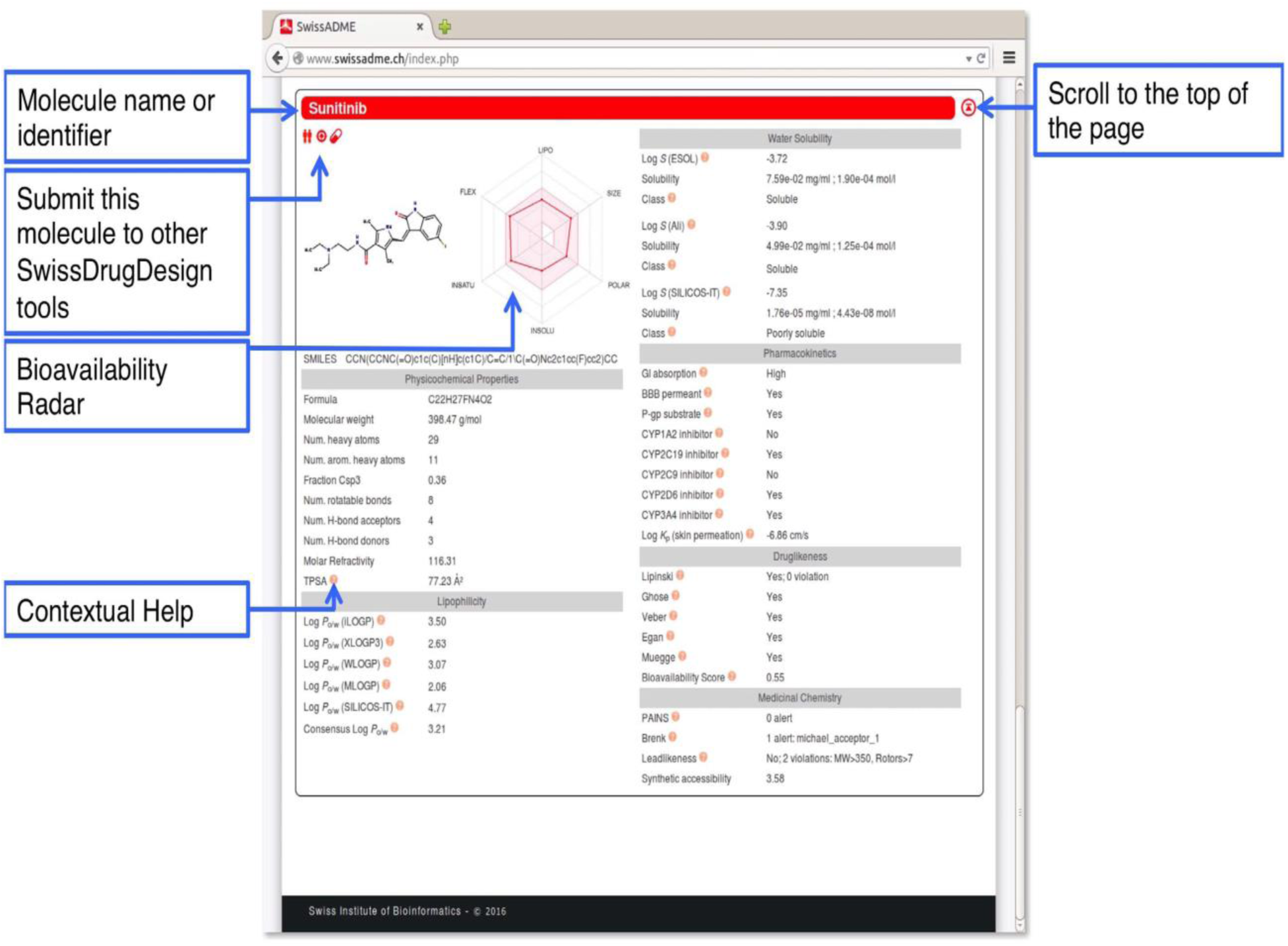
Window showing the outcome in the form of Computed parameter values grouped in the different sections of the one-panel-par-molecule output.

Molecular docking of target receptor proteins (dopamine transporter, serotonin transporter, and monoamine oxidase B) was prepared by retrieving the three-dimensional crystal structure from the Protein Data Bank (https://www.rcsb.org/) with PDBID:4M48 for dopamine transporter, PDBID:6DZV for serotonin transporter, and PDBID:1GOS for monoamine oxidase B, as shown in Figure 3.

**Fig. 3.**
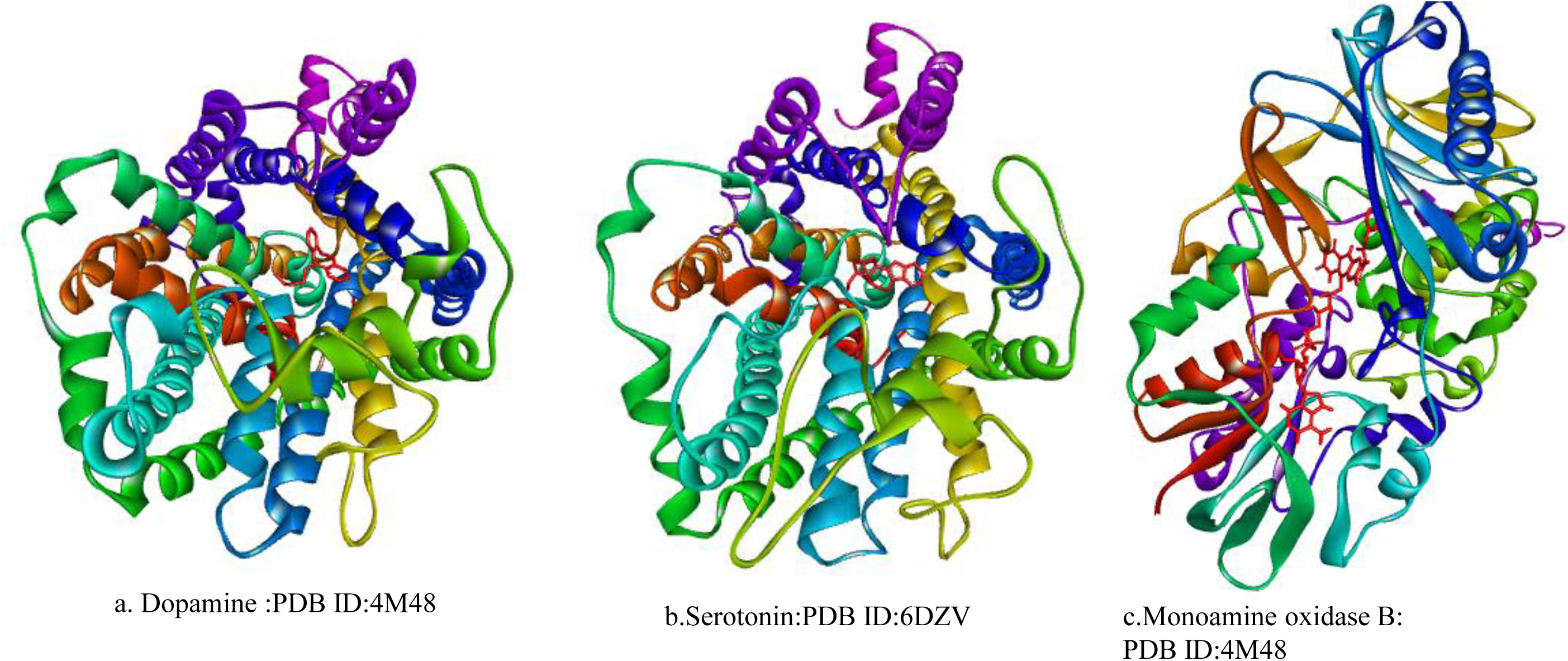
a. Crystal structure of Dopamine complex with Nortriptyline, b. Serotonin complex with ibN-acetyl-D-Glucosamine and, c. Monoamine oxidase b complex with Pargyline

### 2.2 Receptor preparation

The protein was subsequently cleaned by removing the water molecules and any other non-protein atoms and correcting any missing atoms, followed by management of its conformer and the minimization process. The CHARMM27 force field was then applied in the Accelrys Discovery Studio 2.0 software package.

### 2.3 Ligand preparation

Three bioactive main constituents (crocin, picrocrocin, and safranal) were reported in NCBI-PubChem, as shown in the table. 1. These bioactive molecules were selected from PubChem in the SDF format, saved, and converted into PDB structure with the help of Discovery Studio 2.0 software package. Thereafter, by adding charge and torsion, all ligands were saved in PDBQT format. This PDBQT molecule was further used for docking studies.

### 2.4 Molecular Docking Using MGL Tools and AutoDock 4.2

Molecular docking studies were performed using the MGLTools suite and AutoDock 4.2. Input files in PDBQT format were prepared by adding polar hydrogens and assigning Gasteiger charges using AutoDock Tools (version 1.5.6). Following energy minimization, the target protein was placed within a grid box measuring 26.462 Å × 25.591 Å × 22.168 Å along the x, y, and z axes, respectively. The grid spacing was set to 1.00 Å to define the binding site, and all docking parameters were recorded in the AutoDock configuration file for subsequent calculations. Bioactive molecules from saffron, including safranal, crocin, and picrocrocin, were docked into the binding sites of target proteins — specifically, the dopamine transporter, serotonin transporter, and monoamine oxidase B. The docking interactions were compared to those observed in the inhibitor-bound crystal structures of the respective target proteins. The docking results included the binding energy values, reported in kcal/mol.

### 2.5 Analysis and Output Visualization using Discovery Studio

The docking poses obtained were ranked based on their docking scores, which predict the binding affinity of the ligands to the target proteins. The scoring function implemented in AutoDock was used to estimate the binding free energy between the ligand and receptor molecules. Saffron’s bioactive compounds were selected based on their expected ability to pass through the blood-brain barrier and bind. Key interacting amino acid residues within the binding sites were visualized and analyzed using the Discovery Studio software package to better understand ligand–receptor interactions.

### 2.6 Pharmacokinetic and Toxicity evaluation

Bioactive molecules (safranal, crocin, and picrocrocin) of saffron were evaluated for their pharmacokinetic profile and the presence of any toxic effects. The pharmacokinetic profile of a lead molecule is predicted based on Lipinsky’s rule of five and the Veber rule by evaluating their physicochemical properties like calculated partition coefficient (cLogP), molecular weight, and hydrogen bond donor and acceptor sites (Lipinski, 2003;Pollastri, 2010). The physicochemical properties of the bioactive molecules were calculated by using DataWarrior software. The presence of (if any) toxic effects in the screened lead molecules is evaluated from the different functional groups present in the lead molecule. The toxicity evaluation is performed by using Osiris Molecular Property Explorer software. In the Osiris Molecular Property Explorer software, there is a presence of major toxic effects such as mutagenicity, tumorigenicity, irritant effects, and reproductive effects in the lead molecules. This software searches for the presence of those functional groups that are responsible for the toxic effects of the ligand molecule. The physicochemical parameters, as well as the toxicity profile of all the selected ligand molecules, were evaluated by using DataWarrior software. DataWarrior software checks for the presence of major toxic effects such as mutagenicity, tumorigenicity, irritant and reproductive effects in the lead molecules. This software searches for the presence of those functional groups that are responsible for the toxic effects of the ligand molecule. The Data Warrior program also calculates the drug-likeness and drug score of the lead molecule based on their physicochemical properties, such as absorption, distribution, metabolism, excretion, and toxicity.

### 2.7 Blood-Brain barrier permeability

It is important for any drugs or natural bioactive molecules to cross the blood-brain barrier for the treatment of human central nervous system diseases such as schizophrenia, Alzheimer’s, epilepsy, and depression. The SWISS ADME prediction tool is used to generate the Swiss boiled egg compartment, which revealed the blood-brain permeability of safranal, as shown in Fig.4. The safranal molecule also showed a PGP, indicating that safranal is unable to effluate from the central nervous system by the P-glycoprotein. Whereas no blood-brain barrier (BBB) permeability was shown by the crocin and picrocrocin.

**Fig. 4.**
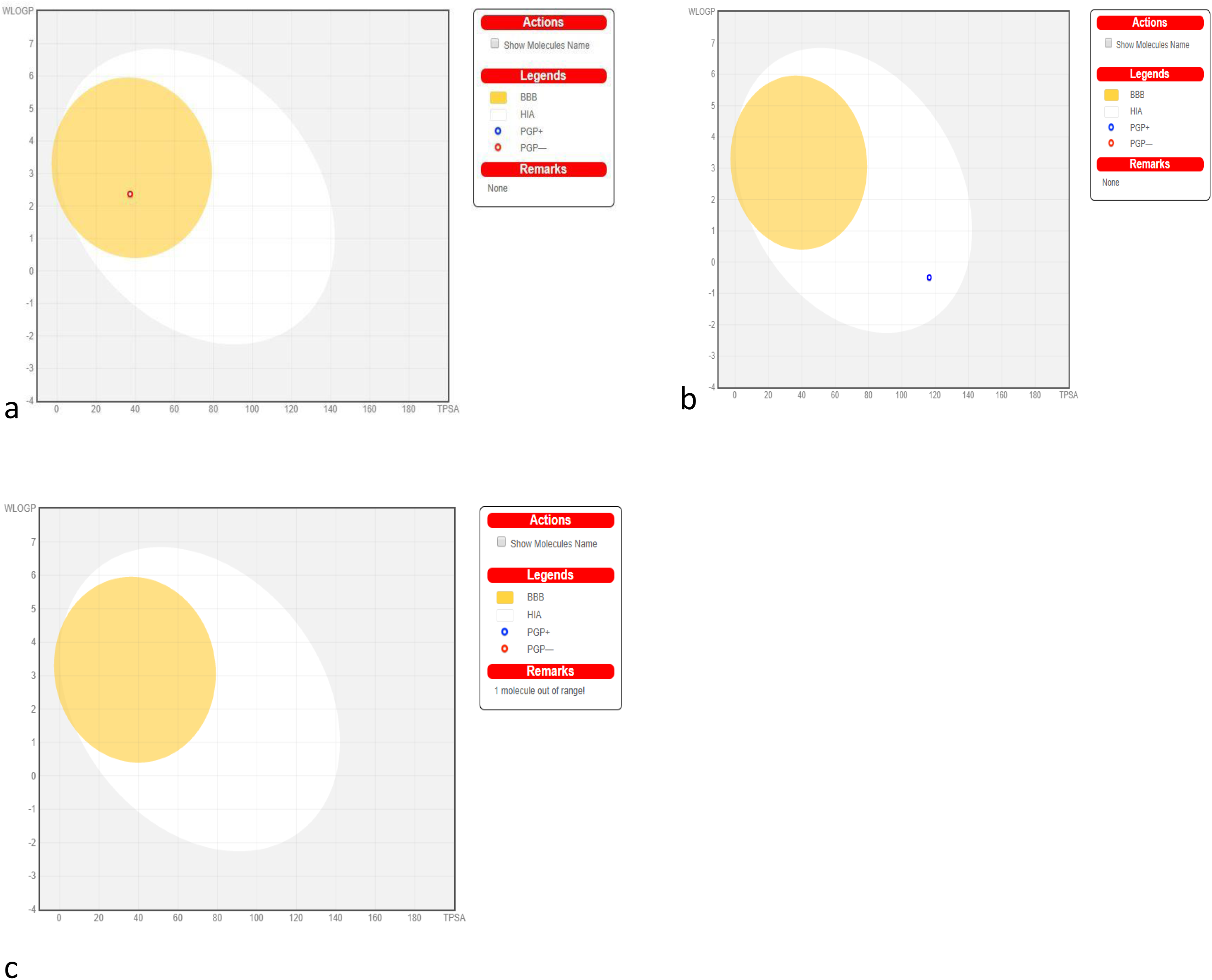
The boiled egg compartment model to graphically represent the blood brain penetration of ligands (Picrocrocin, safranal and crocin). a. Graphical representation of safranal as a dot within yellow yolk are well penetrated within the brain. Red dots represent that safranal could not be the substrate for P-glycoprotein indicating the good absorption and penetration to the brain. While Picrocrocin (b) and Safranal (c) does not represent the dot within the yellow yolk indicate no blood brain permeability

### 2.8 Molecular Docking Validation

The molecular docking process for the docking of bound ligands nortriptyline, N-acetyl-D-glucosamine, and pargyline against dopamine transporter (4M48), Serotonin transporter (6DZV), and monoamine oxidase B, respectively, was performed by considering the binding energy.

### 2.9 Binding Energy

The binding energy obtained by performing molecular docking simulation of the bound ligands Nortriptyline, N-acetyl-D-Glucosamine, and Pargyline against dopamine transporter (4M48) is - 8.38 kcal/mol, serotonin transporter (6DZV) is -9.07 kcal/mol, and monoamine oxidase B (1GOS) is -5.79 kcal/mol, respectively, as shown in Table 2.

## 3. Results

Binding energy and affinity of ligands (Nortriptyline, N-acetyl-D-Glucosamine, and Pargyline) and saffron constituents (crocin, picrocrocin, and safranal) towards the Dopamine transporter (4M48), Serotonin transport protein (6DZV), and Monoamine oxidase B (1GOS) The binding energy calculated from the molecular docking simulation of Nortriptyline with the Dopamine transporter (4M48) is -8.38 Kcal/mol, showing a binding strength of 0.72 µM, as shown in Table 2. Nortriptyline interacts with the amino acid residues ALA A:117, TYR A:124, VAL A:120, ALA A:479, PHE A:43, and PHE A:319 by van der Waals forces. Conventional hydrogen bond, carbon-hydrogen bond, pi-donor hydrogen bond, pi-sigma bond, pi-pi T-shaped bond, alkyl bond, and pi-alkyl bond at the active site of the dopamine transporter. Conversely, several non-interacting amino acid residues, namely ASP A:121, SER A:422, GLY A:425, VAL A:327, PHE A:325, ILE A:116, ILE A:483, GLY A:322, ALA A:44, SER A:421, and ASP A:46, are located at the active site of the dopamine transporter, as illustrated in Fig. 5. Picrocrocin and safranal demonstrate binding energies of -6.51 kcal/mol and -5.20 kcal/mol, respectively, against the dopamine transporter, which falls within the recommended range of -5 to -15 Kcal/mol, with binding affinities of 16.98 µM and 154.87 µM, respectively, as seen in Table 2. The binding energy and binding affinity of crocin were determined to be in an unfavorable range with the dopamine transporter. Picrocrocin was observed to be docked at the active site of the dopamine transporter, interacting with the amino acid residues ASP A:46, PHE A:319, SER A:320, PHE A:43, TYR A:124, VAL A:120, ALA A:117, VAL A:327, and ALA A:44 through van der Waals forces, conventional hydrogen bonds, carbon-hydrogen bonds, alkyl interactions, and pi-alkyl bonds. Conversely, several non-interacting amino acid residues, namely ALA A:48, LEU A:321, PHE A:325, ILE A:116, ILE A:483, GLY A:322, GLY A:425, SER A:421, SER A:422, and ASP A:121, are illustrated in Fig. 8. Safranal interacts with the active site of the dopamine transporter through amino acid residues PHE A:325, VAL A:120, VAL A:327, ALA A:479, PHE A:43, ILE A:483, VAL A:113, ILE A:116, and ALA A:117 via van der Waals forces, conventional hydrogen bonds, carbon-hydrogen bonds, pi-sigma interactions, alkyl bonds, and pi-alkyl bonds. The non-interacting amino acid residues, GLY A:425 and ALA A:428, are present in the docking pose of the dopamine transporter–safranal complex, as illustrated in Fig. 11.

**Fig. 5.**
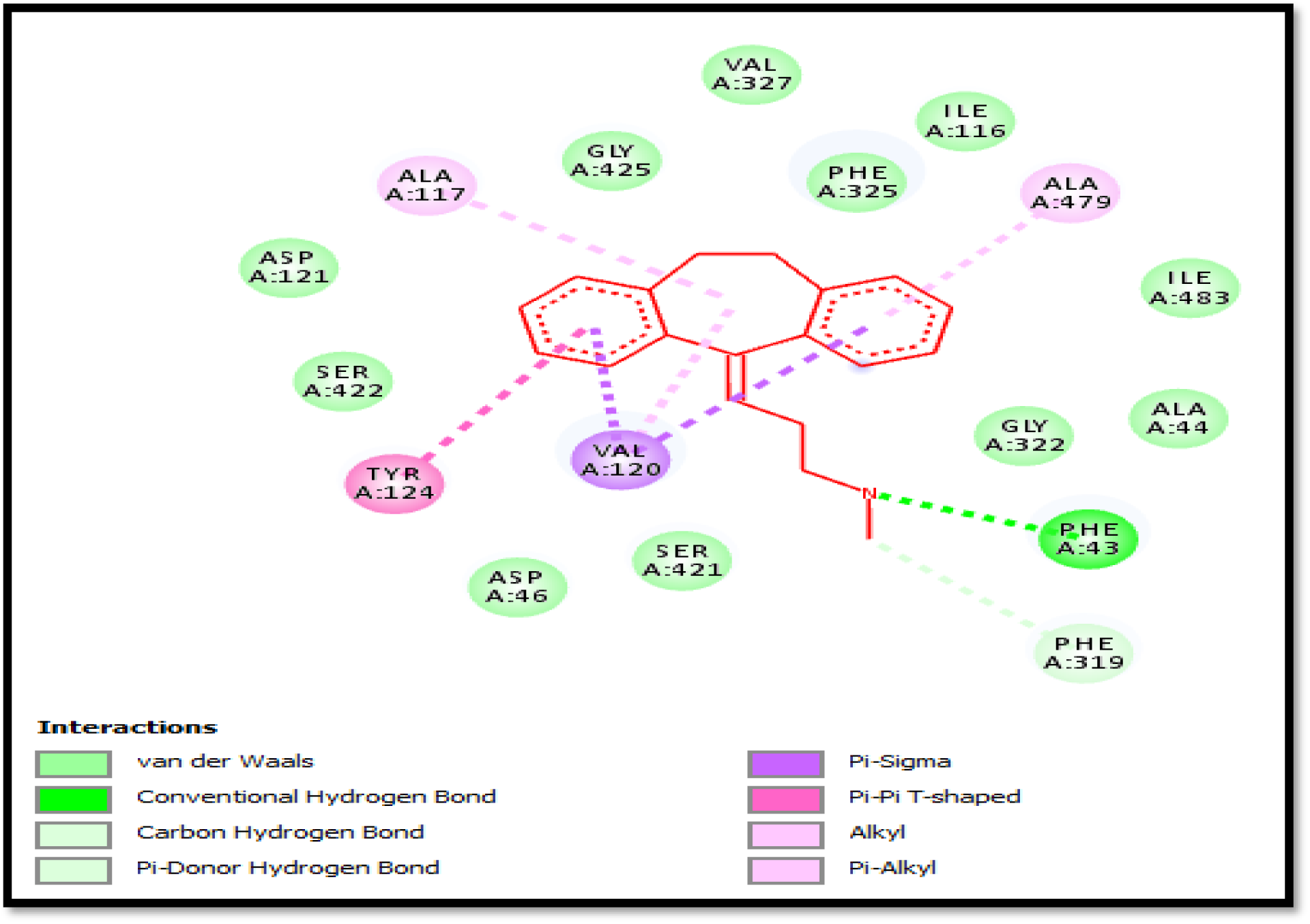
Two-dimensional binding mode and chemical interactions of dopamine with bound ligand (Nortriptyline) complex

N-acetyl-D-glucosamine was observed to be docked at the active site of the serotonin transporter protein (6DZV) with a binding energy of -9.07 kcal/mol and a binding affinity of 0.23 µM, as seen in Table 2. N-acetyl-D-glucosamine interacts with the amino acid residues ALA A:169, GLY A:442, TYR A:95, and PHE A:319 through van der Waals, pi-sigma, pi-pi T-shaped, alkyl, amide-pi-stacked, and pi-alkyl interactions at the active site of the serotonin transporter protein. Conversely, several non-interacting amino acid residues, namely LEU A:443, THR A:439, SER A:438, ASP A:98, SER A:336, LEU A:337, ALA A:96, PHE A:335, GLY A:338, and ILE A:172, are located at the active site of the serotonin transporter receptor, as illustrated in Fig. 6. Picrocrocin and Safranal were observed to dock at the active site of the serotonin transporter protein, exhibiting a binding energy of -7.21 kcal/mol with a binding affinity of 5.20 µM and - 4.82 kcal/mol with a binding affinity of 293.08 µM, respectively, as presented in Table 2. Nonetheless, crocin interacted with the active region of the serotonin transporter protein; however, it exhibited negligible binding energy and affinity. Amino acid residues, specifically TYR A:95, PHE A:335, VAL A:343, ILE A:172, ASP A:98, and TYR A:176, located at the active site of the serotonin transporter protein, are involved in the interaction with crocin. Van der Waals forces, conventional hydrogen bonds, carbon-hydrogen bonds, pi-sigma bonds, alkyl bonds, and pi-alkyl bonds. Conversely, other non-interacting amino acid residues, i.e., VAL A:97, ALA A:96, SER A:336, LEU A:337, GLY A:338, PHE A:341, GLY A:442, LEU A:443, THR A:439, SER A:438, and ALA A:173 are located near the active site of the serotonin transporter protein, as illustrated in Fig.9. Safranal interacts with the active site of the serotonin transporter protein through amino acid residues ALA A:173, TYR A:176, ILE A:172, LEU A:443, and ASN A:177 via van der Waals forces, conventional hydrogen bonds, pi-sigma interactions, alkyl bonds, and pi-alkyl interactions. Some amino acid residues that do not interact, like SER A:438, GLY A:442, PHE A:263, and THR A:439, can also be found in the docking position of the safranal-serotonin transporter protein complex, as shown in Fig. 12. The binding energy from the molecular docking simulation of Pargyline with Monoamine oxidase B (1GOS) is -5.79 Kcal/mol, showing a binding strength of 56.83 µM, as shown in Table 2. Pargyline interacts with the amino acid residues ILE A:198, LEU A:171, CYS A:172, TYR A:326, ILE A:199, PHE A:168, LEU A:167, LEU A:164, and TRP A:119 through Pi-Pi T-shaped, alkyl, and Pi-alkyl interactions at the active site of monoamine oxidase B. Conversely, several non-interacting amino acid residues, namely TYR A:435, GLN A:206, and ILE A:316, are located at the active site of the dopamine transporter, as illustrated in Fig. 7. Picrocrocin exhibited a greater binding energy (-7.63 kcal/mol) compared to the pargyline-bound ligand in the active site of monoamine oxidase B (1GOS), which had a binding affinity of 2.53 µM. Moreover, safranal demonstrates a greater binding energy (-6.11 kcal/mol) compared to the bound ligand in the crystal structure, which has a binding affinity of 3.34 µM, as illustrated in Table 2. The amino acids at the active site of monoamine oxidase B (1GOS), such as GLY A:205, GLN A:65, MET A:436, TYR A:60, TYR A:398, TYR A:435, PHE A:343, TYR A:326, CYS A:172, LEU A:171, and GLN A:206, connect with picrocrocin using alkyl and pi-alkyl bonds. Conversely, some non-interacting amino acid residues, namely GLY A:62, VAL A:61, GLU A:437, GLY A:58, LEU A:328, and MET A:341, are observed in the docking position of the monoamine oxidase B–picrocrocin complex, as illustrated in Fig. 10. Safranal interacts with the active site of monoamine oxidase B by using the amino acid residues TYR A:435, MET A:436, and GLN A:206 through alkyl and pi-alkyl bonds. The non-interacting amino acid residues, specifically GLU A:437, VAL A:61, GLN A:65, TYR A:60, GLY A:205, SER A:59, and GLY A:58, are present in the docking position of the monoamine oxidase B–safranal complex, as illustrated in Fig. 13.

**Fig. 6.**
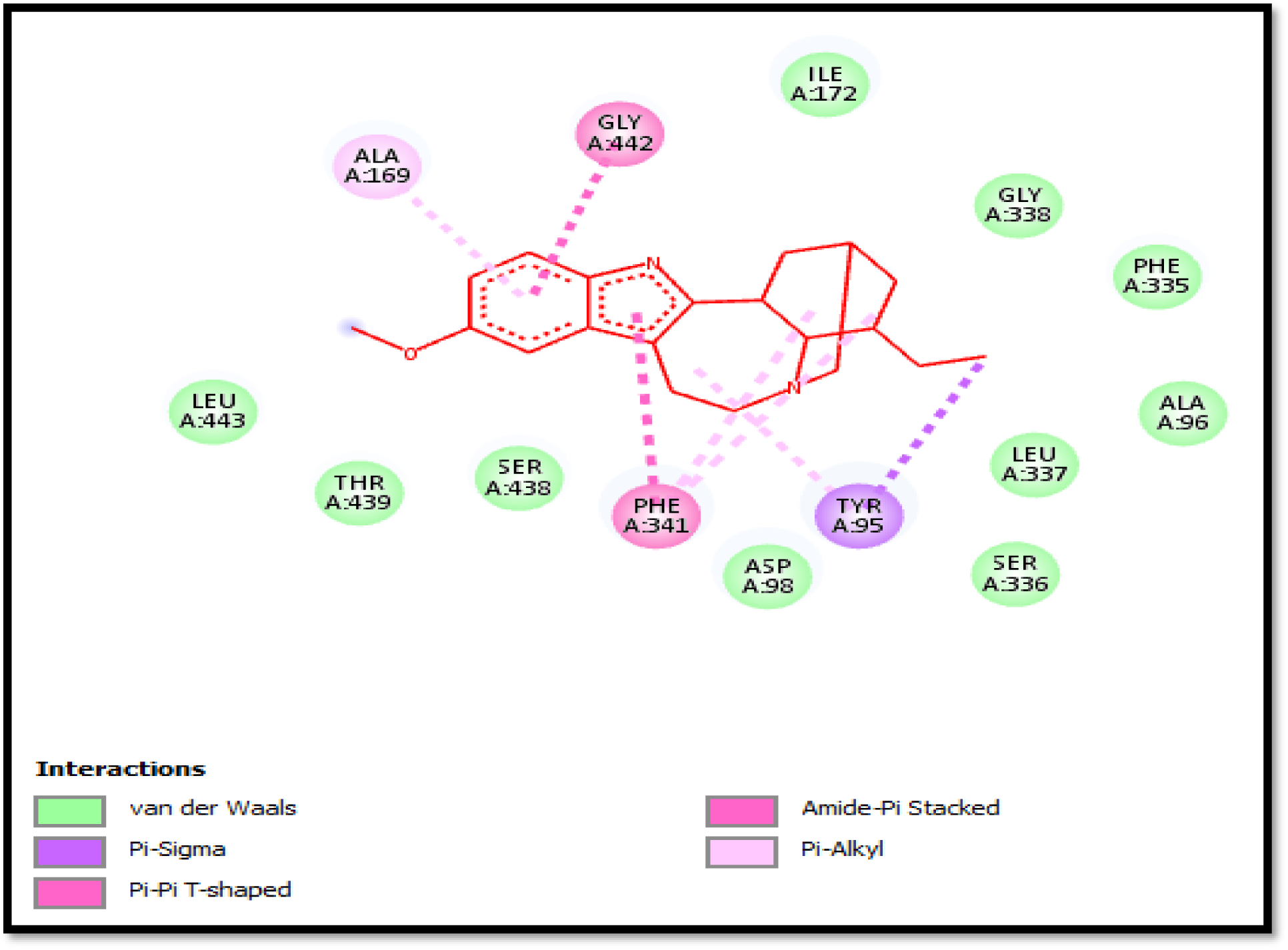
Two-dimensional binding mode and chemical interactions of Serotonin with bound ligand (N-acetyl-D-Glucosamine) complex

**Fig. 7.**
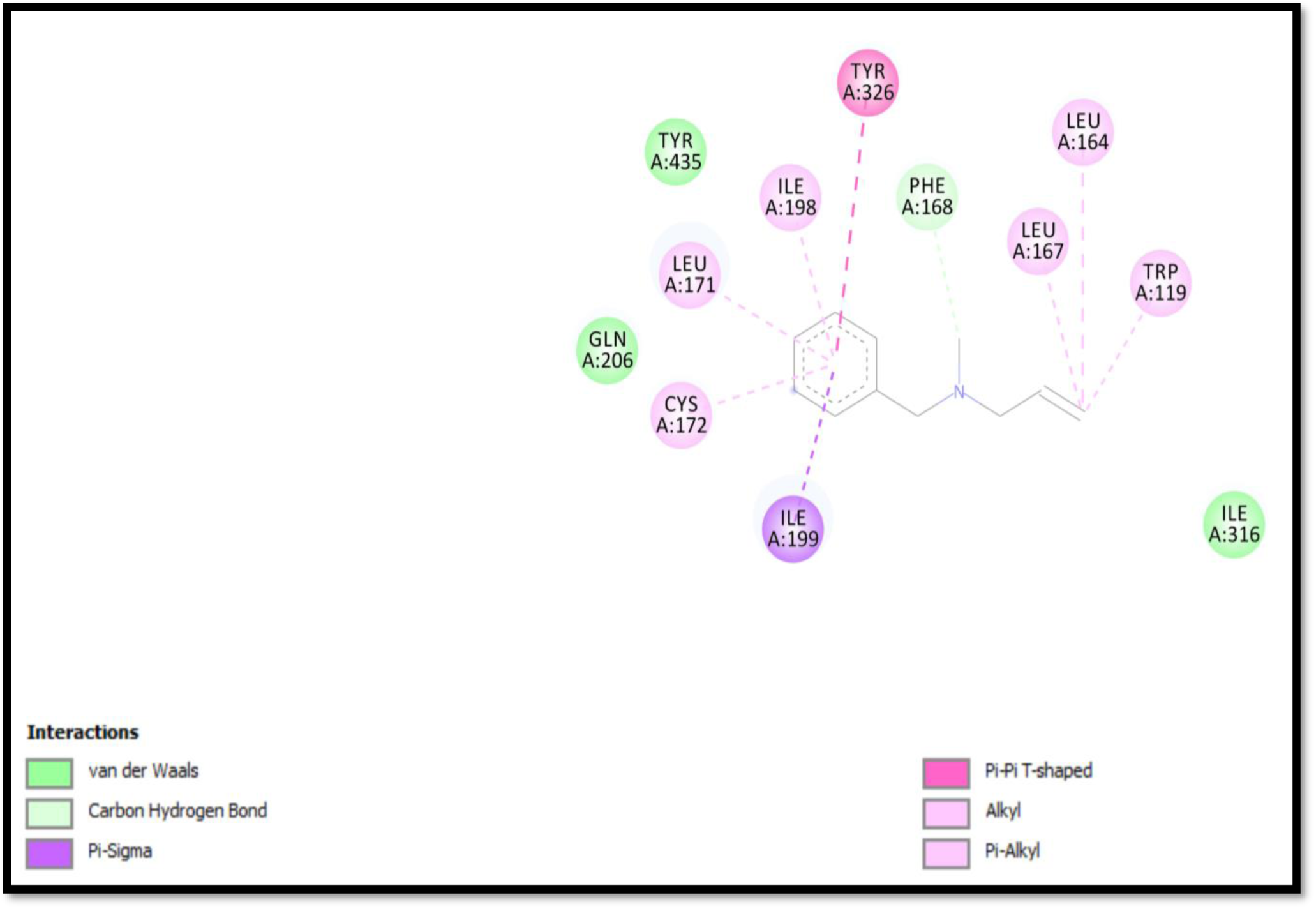
Two-dimensional binding mode and chemical interactions of Monoamine oxidase b with bound ligand (Pargyline) complex

**Fig. 8.**
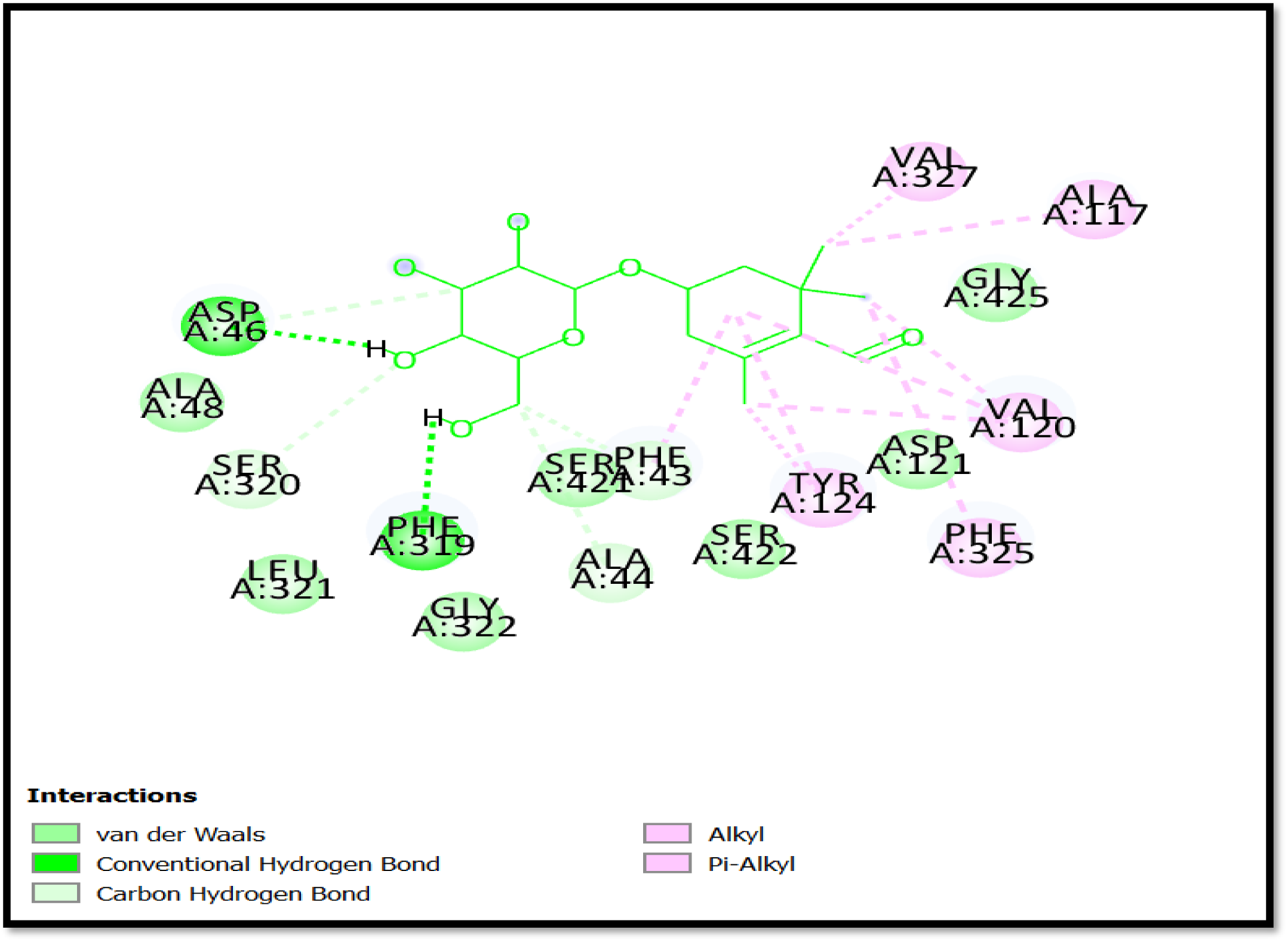
Two-dimensional binding mode and chemical interactions of Picrocrocin with Dopamine transporter protein

**Fig. 9.**
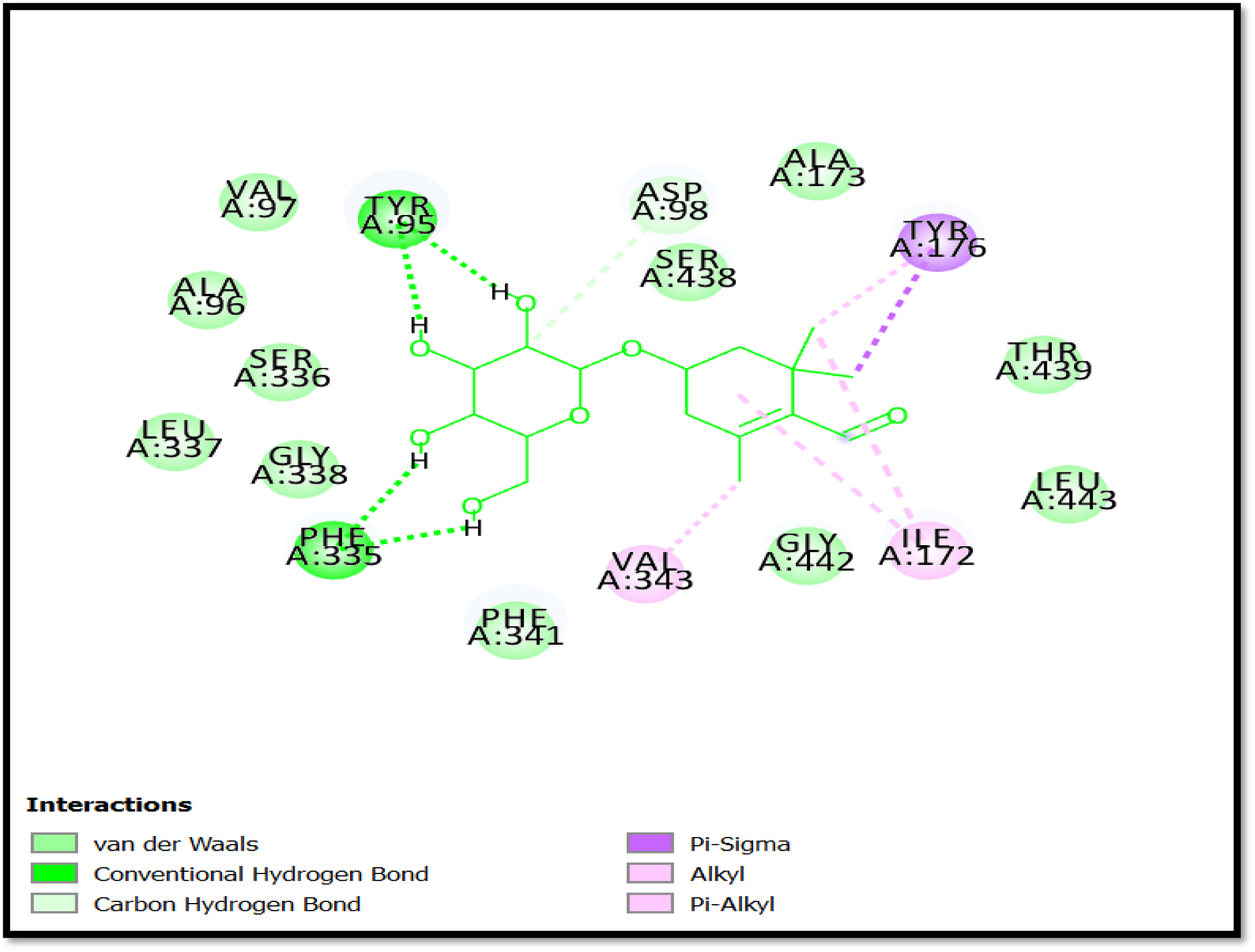
Two-dimensional binding mode and chemical interactions of Picrocrocin with Serotonin transporter protein

**Fig. 10.**
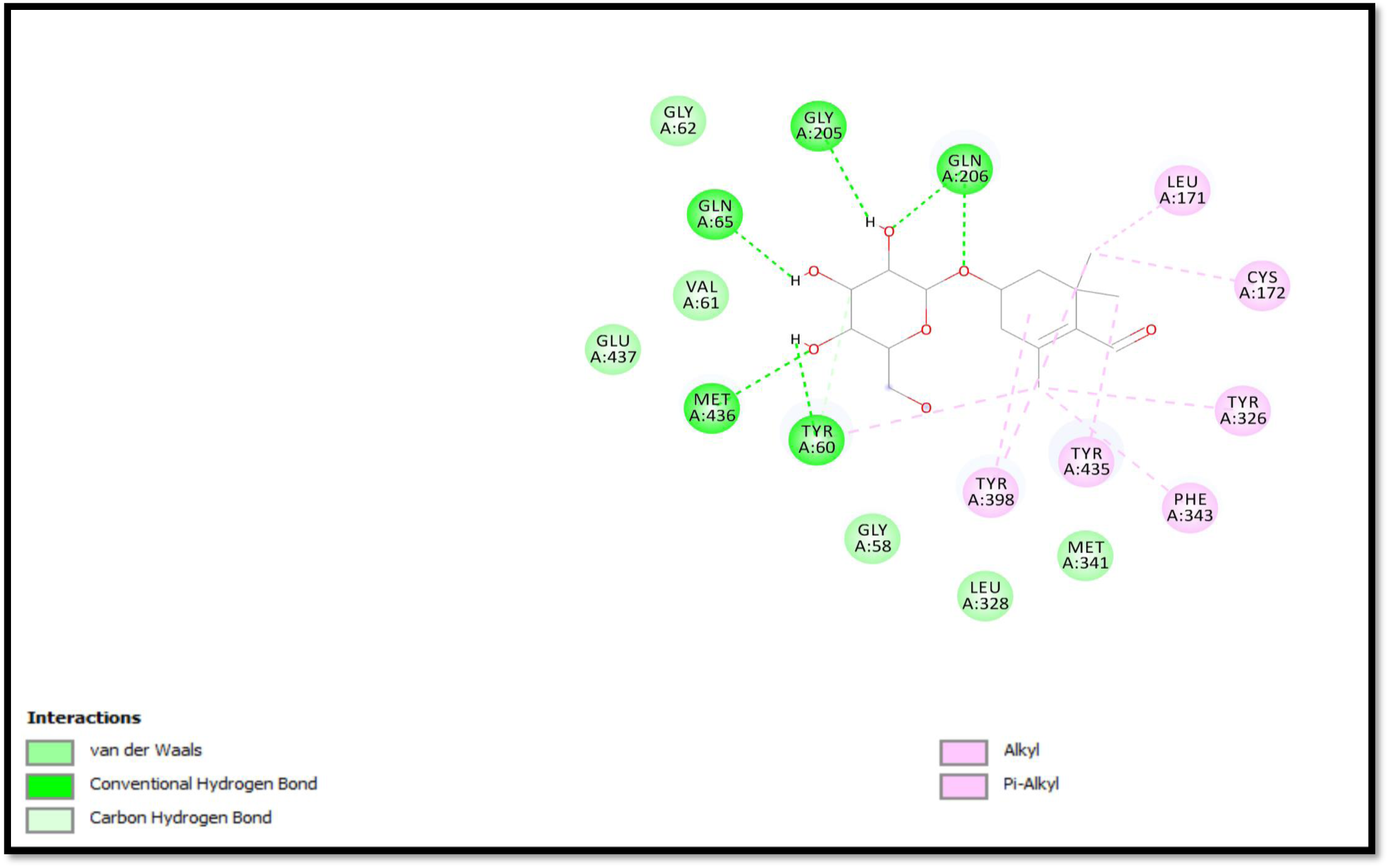
Two-dimensional binding mode and chemical interactions of Picrocrocin with Monoamine oxidase b.

**Fig. 11.**
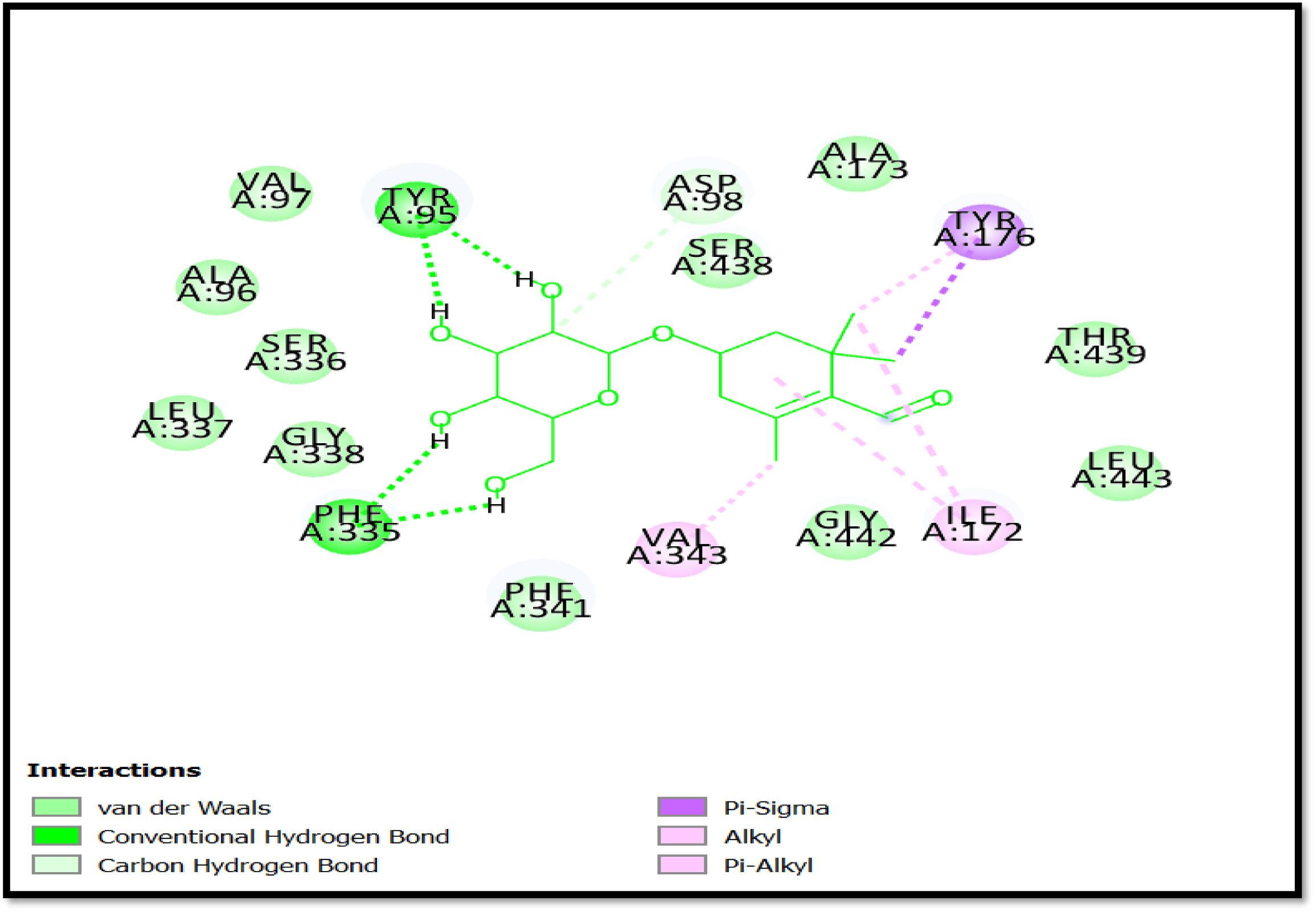
Two-dimensional binding mode and chemical interactions of safranal with Dopamine transporter protein

**Fig. 12.**
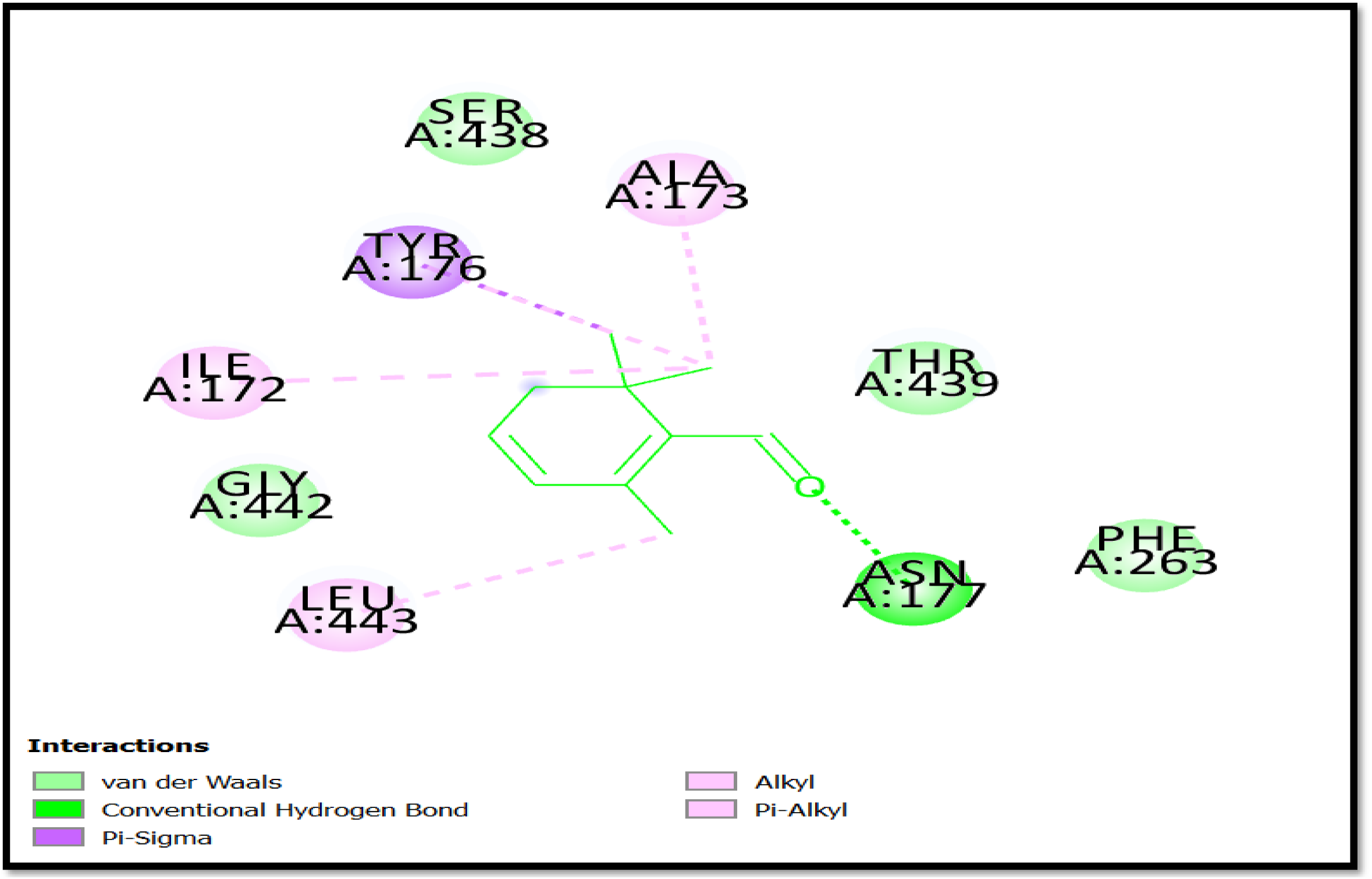
Two-dimensional binding mode and chemical interactions of safranal with Serotonin transporter protein

**Fig. 13.**
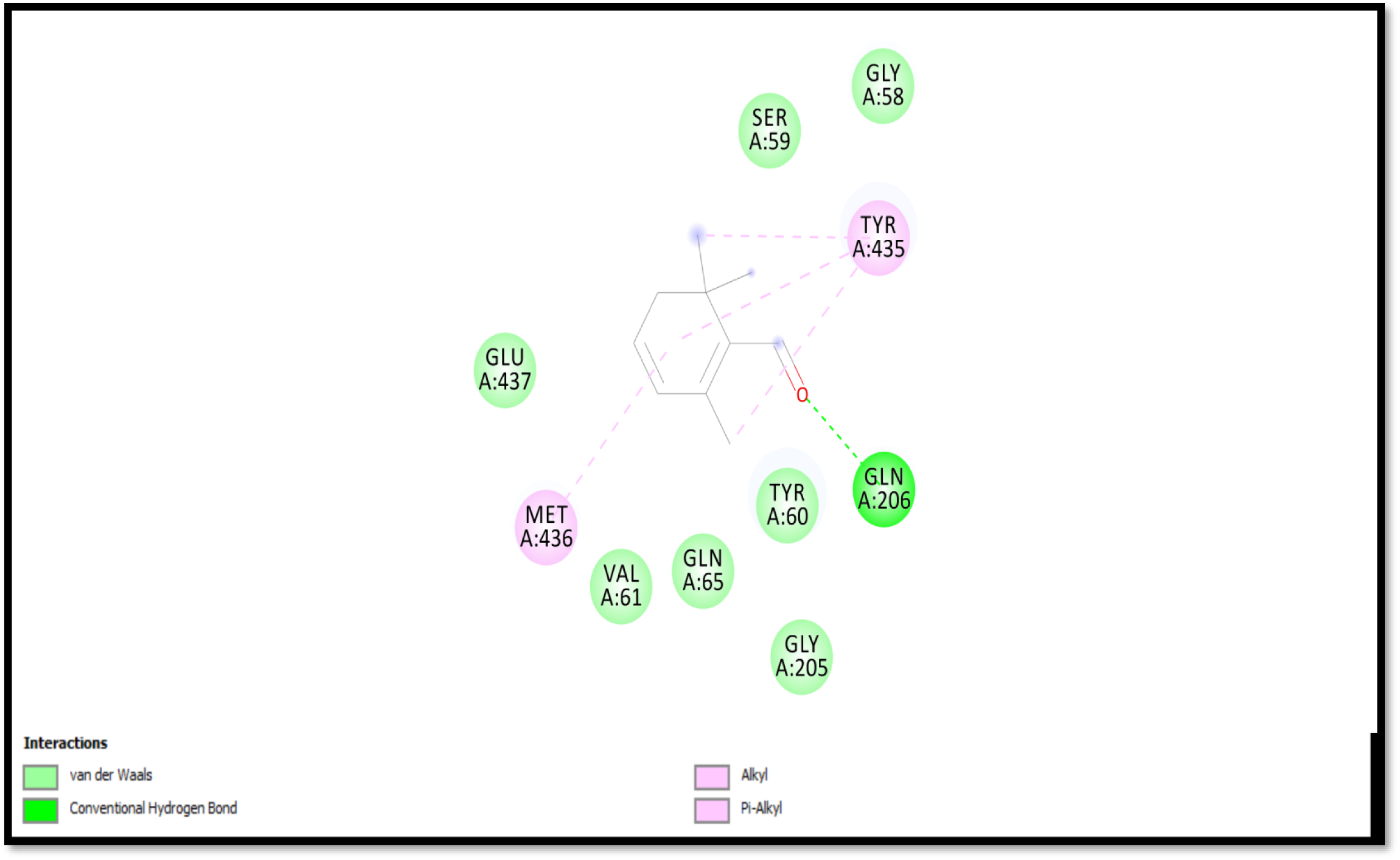
Two-dimensional binding mode and chemical interactions of safranal with Monoamine oxidase b.

### Assessment of pharmacokinetics and toxicity

The ADME and toxicity data of the bioactive compound are presented in Table 3, respectively. The molecular weight of crocin (976 Dalton) exceeds the optimal range for medicinal molecular weight (<500). Safranal (150D) and picrocrocin (330D) possess a suitable molecular weight for optimal drug-likeness. The selected bioactive compounds of saffron exhibited non-mutagenicity, non-tumorigenicity, and no reproductive effects, as determined using Data Warrior software. However, picrocrocin and safranal have irritating properties, unlike crocin. The lipophilicity, measured by cLogP (calculated partition coefficient), of bioactive compounds such as safranal (1.84), crocin (-1.9), and picrocrocin (-0.57) falls within an acceptable range of -4 to 5.6. The topological polar surface area (TPSA) of picrocrocin and safranal, as presented in Table 3, was determined to be within the permitted range (<140 Å).

## 4. Discussion

Saffron components possess antioxidant and anti-inflammatory properties, as well as therapeutic promise for many neuropsychiatric disorders, including Parkinson’s disease and Alzheimer’s disease (El Midaoui et al., 2022). Numerous human and animal studies have demonstrated the advantageous effects of saffron components in treating depression by enhancing serotonin availability to presynaptic neurons (Shafiee et al., 2025). Ettehadi et al. (2013) demonstrated an increase in dopamine neurotransmitter levels in the brain following the injection of *C. sativus* extract in rats, indicating the interaction of saffron components with the dopaminergic system. The elevation of dopamine and serotonin concentrations in the brain is also slightly concerning for the enzyme monoamine oxidase B, which plays a crucial role in regulating dopamine and serotonin levels (Juárez Olguín et al., 2016; Azizi, 2022). Moreover, research indicates that the inhibition of monoamine oxidase B can exhibit antidepressant effects in individuals with depression by prolonging the availability of serotonin and dopamine for effective synaptic transmission (Suchting et al., 2021;Laban and Saadabadi, 2023). Furthermore, the pretreatment of saffron tea in mice exposed to aflatoxin B1 mitigates learning and memory deficits by reducing the activity of monoamine oxidase isoforms A and B (Linardaki et al., 2017). Consequently, the investigations demonstrate that saffron compounds interact with many components of the neurotransmitter system, including the serotonin transport protein, dopamine transporter protein, and monoamine oxidase B. Consequently, we have examined the interaction between saffron compounds and neurotransmitter system components by in silico molecular docking. A significant characteristic of the central nervous system is its separation from the bloodstream by the blood-brain barrier. The blood-brain barrier (BBB) primarily comprises tight junctions formed by endothelial cells of brain capillaries, astroglial cells, perivascular macrophages, pericytes, and the basal lamina, which effectively limits the passage of molecules from the bloodstream into the brain, allowing only small and lipophilic substances essential for brain function, such as cofactors, vital nutrients, and precursors (Abbott et al., 2010; Upadhyay, 2014; 2014;Greene et al., 2019; Gawdi et al., 2023). Consequently, any neurotherapeutic drugs must possess a crucial attribute that enables them to traverse the blood-brain barrier and access the central nervous system. In this study, we examine the blood-brain barrier permeability of saffron compounds (picrocrocin, safranal, and crocin) utilizing the in silico SWISS ADME web server. Among the elements of saffron, safranal has demonstrated blood-brain barrier permeability, as illustrated in Fig. 5a. Research has indicated that the permeability of neurotherapeutic agents across the blood-brain barrier is insufficient to provide the desired therapeutic effects; these compounds must also possess the capability to persist in the brain for an extended duration. P-glycoprotein is a type of protein found on the surface of blood vessel cells that helps move certain drugs out of the brain, making it harder for them to get in. But safranal stays in the brain because these proteins don’t move it out. Nonetheless, safranal has demonstrated that it is not a substrate of P-glycoprotein transporters, as indicated in Fig. 5a, which prevents the efflux of safranal from the brain. It might be posited that safranal is the more potent component of saffron, capable of traversing the blood-brain barrier to enter the central nervous system and elicit a desired effect. Additionally, the Lipinski rule of five states that for a drug to be effectively absorbed by cells, it must have certain physical and chemical properties, such as being fat-friendly (0.4-5.6), having a topological polar surface area (TPSA) greater than 140 Å, a molecular weight over 500 Da, and similarity to other drugs. Picrocrocin and safranal have acceptable pharmacokinetic features, as seen in Table 3. However, the molecular weight of crocin was determined to be 976 Daltons, exceeding the permissible limit of 500, which adversely affects its bioavailability. Moreover, picrocrocin, safranal, and crocin have non-mutagenic, non-tumorigenic, and non-reproductive effects. Consequently, the blood-brain barrier permeability, pharmacokinetic characteristics, and toxicity assessment suggest the efficacy of safranal as a drug-like chemical.

Dopamine is a neurotransmitter released by dopaminergic neurons primarily located in the substantia nigra, ventral tegmental area (VTA) of the midbrain, corpus striatum, and nucleus accumbens within the central nervous system. It plays a crucial role in dopaminergic transmission related to behavior, problem-solving, decision-making, sensory perception, feelings of happiness and pleasure, food intake, and social functioning (Salatino-Oliveira et al., 2018; Cinque et al., 2018). Abnormal dopaminergic transmission has been observed to reflect the pathophysiology underlying major depressive disorder and other neurocognitive functions, including anhedonia, diminished motivation and drive, and reduced physical energy (Brigitta, 2002; Epstein and Silbersweig, 2015; Belujon and Grace, 2017). Research has demonstrated that patients with severe depressive disorders exhibit elevated expression of the dopamine transporter (DAT) at presynaptic neurons in specific brain regions, such as the striatum (Juárez Olguín et al., 2016; Pizzagalli et al., 2019). The physiological function of DAT is to terminate dopaminergic transmission by reabsorbing dopamine from the synaptic cleft into presynaptic neurons, where it is subsequently repackaged into tiny vesicles for re-release (Banerjee and Lidsky, 1990). Furthermore, an increased level of DAT resulted in a diminished binding availability of dopamine to dopamine receptors, highlighting its crucial role in the pathophysiology of depression (Kugaya et al., 2003). Bupropion is a dopamine transporter inhibitor employed as an antidepressant to elevate synaptic dopamine levels by obstructing dopamine reuptake (Stahl et al., 2004). Research indicates that saffron ingredients exert an antidepressant effect by regulating neurotransmitter levels (dopamine and serotonin) in the brain (Wang et al., 2010; 2010;Lopresti and Drummond, 2014). Consequently, we have analyzed the molecular docking studies of saffron ingredients (picrocrocin, safranal, and crocin) against the dopamine transporter (4M48), serotonin transport protein (6DZV), and monoamine oxidase B (1GOS). We determined the notable binding energy and binding affinity of picrocrocin (-6.51 kcal/mol; 16.98 µM) and safranal (-5.20 kcal/mol; 154.87 µM) for the dopamine transporter (4M48). Picrocrocin and safranal interact similarly with the amino acids ALA A:117, TYR A:124, VAL A:120, ALA A:479, PHE A:43, and PHE A:319 that are part of the crystal structure of the dopamine transporter complex with nortriptyline. Likewise, in relation to the serotonin transport protein (6DZV), Picrocrocin (-7.21 Kcal/mol, 5.20 µM) and safranal (-4.82 Kcal/mol, 293.08 µM) demonstrate notable binding energy and binding affinity, respectively. Picrocrocin exhibits analogous interactive amino acid residue TYR A:95 and non-interacting amino acid residues LEU A:337, GLY A:338, and PHE A:341, as shown in the crystal structure of the serotonin transport protein complexed with N-acetyl-D-glucosamine. The binding energies of picrocrocin and safranal to monoamine oxidase B are stronger than those of the pargyline ligand in the crystal structure of monoamine oxidase B, with binding strengths of 2.53 µM and 3.34 µM, respectively. Compared to pargyline, picrocrocin has similar amino acids that interact with monoamine oxidase B, which are TYR A:326, LEU A:171, and CYS A:172, and also has some that do not interact, like TYR A:435 and GLN A:206, as shown in Fig. 10. GLN A:206, a non-interacting amino acid located at the docking position of safranal, exhibits similarities to the docking configuration of the pargyline-monoamine oxidase B complex, as illustrated in Fig. 13. Among the components of saffron, picrocrocin and safranal show a strong ability to bind, good properties for how they move in the body, and important interactions with the dopamine transporter (4M48), serotonin transporter protein (6DZV), and monoamine oxidase B (1GOS), suggesting they could be useful as natural treatments. This action may optimize dopamine and serotonin levels in the synaptic cleft of patients suffering from depression. Additionally, safranal was found to be more effective at blocking the dopamine transporter protein because it can easily cross the blood-brain barrier, has a strong ability to bind, and shares similar interacting parts with how nortriptyline interacts with the dopamine transporter protein.

## 5. Data availability statement

Not applicable

## 6. Ethics statement

Not applicable

## 7. Author contributions

Dr. Brijendra Singh: Conceptualization, Methodology Design, Experimental Investigation, Advanced Data Analysis, Curated Dataset Generation, Bioactive Component Selection Rationale, Manuscript Drafting, Project Leadership.

Dr. Deepak Sharma: Botanical Expertise (Plant Species Selection and Ethnobotanical Rationale), Critical Manuscript Review and Scientific Validation.

Dr. Vyas Madhavrao Shingatgeri: Validation, Manuscript Review and Critical Evaluation, Contribution to Safety and Risk Assessment.

Dr. Vinay Lomash: Research Oversight, Procedural Validation and Quality Assurance, Integrated Efficacy and Toxicology Assessment, Bioactive Component Selection Rationale, Final Manuscript Preparation for Publication.

## 9. Acknowledgments

Not applicable

